# Chitinase 3-like-1 (CHI3L1) in the Pathogenesis of Epidermal Growth Factor Receptor Mutant Non-Small Cell Lung Cancer

**DOI:** 10.1101/2023.09.21.558861

**Authors:** Suchitra Kamle, Bing Ma, Gail Schor, Madison Bailey, Brianna Pham, Inyoung Cho, Hina Khan, Christopher Azzoli, Mara Hofstetter, Chang-Min Lee, Roy Herbst, Katerina Politi, Chun Geun Lee, Jack A. Elias

## Abstract

Non-small cell lung cancer (NSCLC) accounts for 85% of all lung cancers. In NSCLC, 10-20% of Caucasian patients and 30-50% of Asian patients have tumors with activating mutations in the Epidermal Growth Factor Receptor (*EGFR*). A high percentage of these patients exhibit favorable responses to treatment with tyrosine kinase inhibitors (TKI). Unfortunately, a majority of these patients develop therapeutic resistance with progression free survival lasting 9-18 months. The mechanisms that underlie the tumorigenic effects of *EGFR* and the ability of NSCLC to develop resistance to TKI therapies, however, are poorly understood. Here we demonstrate that CHI3L1 is produced by EGFR activation of normal epithelial cells, transformed epithelial cells with wild type *EGFR* and cells with cancer-associated, activating *EGFR* mutations. We also demonstrate that CHI3L1 auto-induces itself and feeds back to stimulate EGFR and its ligands. Highly specific antibodies against CHI3L1 (anti-CHI3L1/FRG) and TKI, individually and in combination, abrogated the effects of EGFR activation on CHI3L1 and the ability of CHI3L1 to stimulate the EGFR axis. Anti-CHI3L1 also interacted with osimertinib to reverse TKI therapeutic resistance and induce tumor cell death and inhibit pulmonary metastasis while stimulating tumor suppressor genes including *KEAP1*. CHI3L1 is a downstream target of EGFR that feeds back to stimulate and activate the EGFR axis. Anti-CHI3L1 is an exciting potential therapeutic for *EGFR* mutant NSCLC, alone and in combination with osimertinib or other TKIs.

## Introduction

Lung cancer is the leading cause of cancer death worldwide (1). Non-small cell lung cancer (NSCLC) is the most common histologic subtype accounting for 85% of all lung cancers (1). Unfortunately, approximately 50% of NSCLCs are advanced enough at diagnosis to preclude successful surgical therapeutic intervention (2). Genomic studies of NSCLC have demonstrated the importance of oncogenic driver mutations in approximately 50% of these cancers with mutations in the epidermal growth factor receptor (*EGFR*) and *KRAS* being most common (3). *EGFR* activating mutations are found in 10-20% of NSCLCs in patients with Caucasian backgrounds and 30-50% of patients with Asian heritage (4). However, the mechanisms that underlie NSCLC in these circumstances are poorly understood.

EGFR (also called *ErbB1* and *Her1*) is a transmembrane receptor tyrosine kinase (TK) that transduces signals that drive cell proliferation, differentiation and motility when the wild type (WT) EGFR binds to its physiologic ligands including epidermal growth factor (EGF), transforming growth factor alpha (TGF-α), amphiregulin and epiregulin (5). EGFR and other *Erb1* family members homo- or hetero-dimerize to activate their tyrosine kinase domains and initiate cellular signaling (5). Although the prevalence of different mutations in EGFR mutant NSCLC can vary depending on the population, the L858R point mutation, exon 19 deletion mutation (41%) and T790M drug resistance mutation are most common. Many patients also have more than one genetic alteration with T790M mutations occurring with L858R or exon 19 deletion alterations (6). These and other mutations cause exaggerated EGFR TK activation with the L858R mutation resulting in an *EGFR* that is significantly more active than its WT control (6). This results in ligand-independent receptor activation which leads to oncogenesis. It also led to the development of therapeutic interventions that target the ATP binding cleft of the EGFR kinase domain (7). There are presently 3 generations of approved tyrosine kinase inhibitor (TKI) therapies (6). Most patients with *EGFR* mutant NSCLC initially respond well to these interventions. However, despite prolonged disease control and high tumor response rates, virtually all patients with *EGFR*-mutant NSCLC eventually develop TKI resistance and manifest tumor progression (8, 9). However, the mechanisms that underlie tumor development, initial TKI responsiveness and the development of TKI therapeutic resistance have not been adequately defined.

Chitinase 3-like-1 (CHI3L1), which is also called YKL-40, is a member of the 18 glycosyl hydrolase gene family where it is the prototypic chitinase-like protein (10). It is produced by a wide variety of cells including epithelial cells, macrophages and tumor cells and functions as a pleiotropic mediator that inhibits cell death and innate immunity while stimulating adaptive type 2 immune responses, M2 macrophage differentiation and fibroproliferative tissue repair (10–15). It is stimulated by a large number of cytokines and other stimuli in the context of injury, inflammation, tissue remodeling and cancer where it can be thought of as part of a fundamental healing response that inhibits injury and apoptosis while driving fibrotic repair (11–13, 15–23). CHI3L1/YKL-40 has nuclear, cytoplasmic, extracellular, and circulating compartments, the latter of which is a biomarker that correlates with disease development and severity (10, 17, 20, 21, 23, 24). Recent studies from our laboratory and others have demonstrated that CHI3L1 has critical roles in neoplasia where it inhibits tumor cell death, inhibits tumor suppressors like p53 and PTEN, stimulates the *BRAF* protooncogene and induces tumorigenic type 2 immune responses and M2 macrophage accumulation (11, 25–27). Most recently it has been appreciated to be a master regulator of anti-tumor immune responses where it stimulates inhibitory immune checkpoint molecules including programmed cell death protein-1 (PD-1) and its ligands PD-L1 and PD-L2, cytotoxic T cell lymphocyte antigen-4 (CTLA4) and its ligands B7.1 and B7.2 and lymphocyte activation gene-3 (LAG3) and suppresses T cell co-stimulation via Icos and Icos ligand (ICOSL) and the CD28 and B7.1 and B7.2 axis (25, 28). Surprisingly, the regulation of CHI3L1 by *EGFR* activation and the role(s) of CHI3L1 in the pathogenesis of *EGFR*-mutant NSCLC have not been investigated. In addition, the role(s) of CHI3L1 in the development of TKI resistance in patients with *EGFR*-mutant NSCLC has not been addressed.

We hypothesized that CHI3L1 plays an essential role(s) in the pathogenesis of the exaggerated activation of EGFR that is seen in *EGFR-*mutation driven NSCLC. We also hypothesized that CHI3L1 is an important mediator in the development of therapeutic resistance in patients with NSCLC on TKI inhibitors such as gefitinib and osimertinib. To test these hypotheses, we defined the effects of EGFR activation on CHI3L1 and the effects of CHI3L1 on EGFR and its physiologic ligands. We also evaluated the effects of TKI inhibitors on the production of CHI3L1, the effects of anti-CHI3L1 and TKI, alone and in combination, on tumor progression and metastasis and the effects of anti-CHI3L1 on therapeutic resistance to osimertinib. These studies demonstrate that EGFR activation stimulates the production of CHI3L1, that CHI3L1 feeds back to form a positive feedback loop by stimulating the production and expression of EGFR and its ligands and that this feedback is abrogated by TKI-based interventions. They also demonstrate that anti-CHI3L1 and TKIs inhibit *EGFR* activation and tumor metastasis, alone and more powerfully in combination. Lastly, they demonstrate that anti-CHI3L1 reverses therapeutic resistance that is seen during chronic osimertinib treatment via a mechanism that enhances the expression of *KEAP1* and other tumor suppressors. These studies demonstrate that CHI3L1 is a critical therapeutic target, alone or in combination with TKIs, in the initial therapy and or salvage therapy of *EGFR*-mutant NSCLC.

## Results

### CHI3L1 is induced by physiologic EGFR activation

Studies were undertaken to define the relationship between CHI3L1 and the EGFR Axis. These investigations were initially undertaken to determine if EGFR activation altered the expression and production of CHI3L1. In these experiments we compared the levels of mRNA encoding CHI3L1 and the accumulation of CHI3L1 protein (YKL-40) in normal human bronchial epithelial cells (NHBE cells) incubated with medium alone or the physiologic ligands of EGFR. EGFR activation of normal epithelial cells by its physiologic ligands EGF and TGF-α stimulated the accumulation of CHI3L1 mRNA and protein (Figure 1, A-C and Supplemental Figure S1). The induction of CHI3L1 mRNA was appreciated after 24 hours and was more prominent after 48-72 hours of incubation (Figure 1A). Similar responses were seen with transformed A549 epithelial cells which have wild-type EGFR. (Figure 1, D-F) These studies demonstrate that the activation of EGFR by its physiologic ligands stimulates CHI3L1 mRNA and protein accumulation.

**Figure 1.**
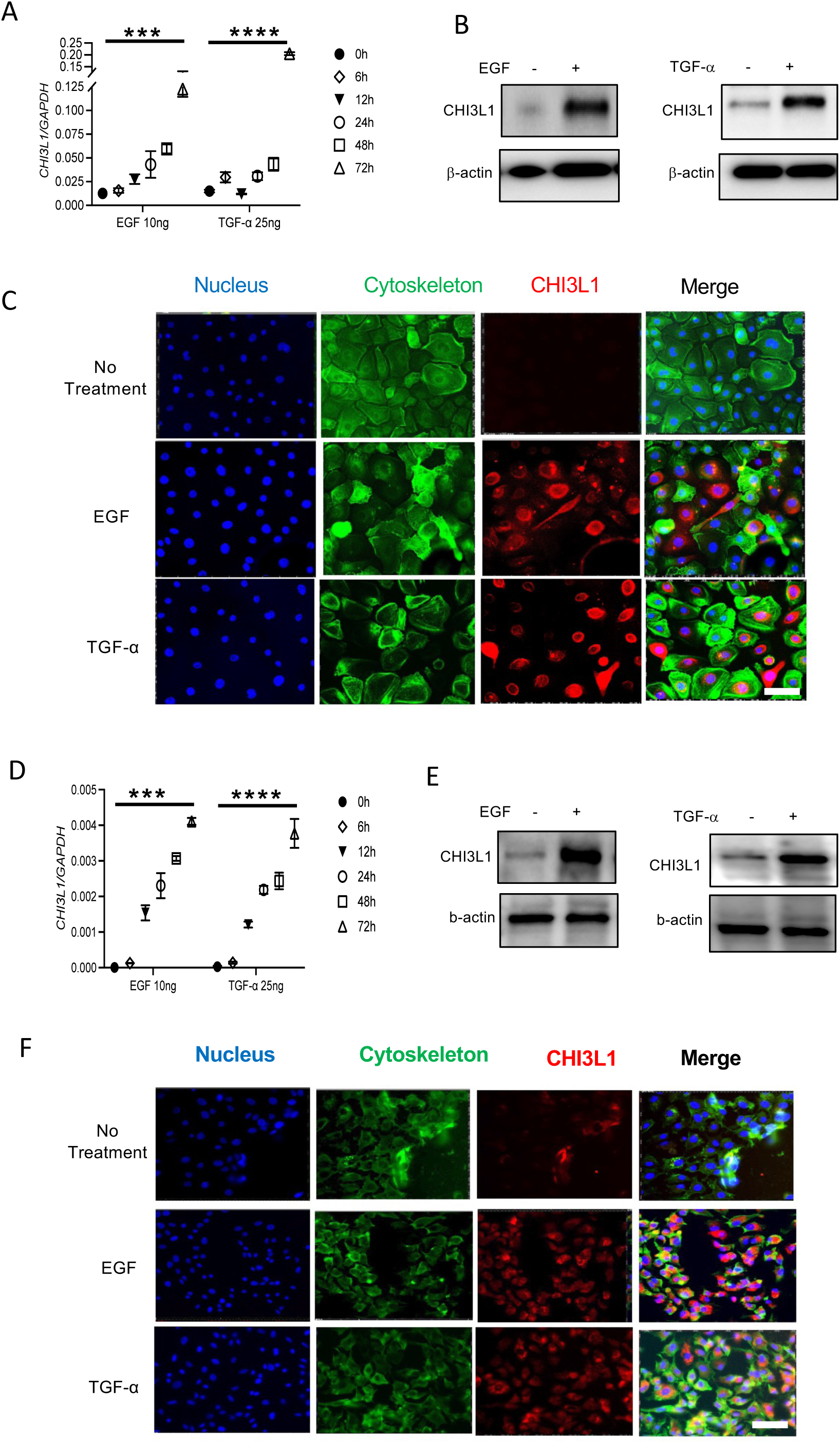
EGFR activation induces the expression and production of CHI3L1. NHBE cells were incubated in the presence or absence of EGF or TGF-α. Triplicate determinations of the levels of mRNA encoding CHI3L1 at the noted time points can be seen in panel A. The accumulation of CHI3L1 protein at 72 hours can be seen with the Western blotting in panel B and IF staining can be seen in panel C. In panels D-F, similar evaluations were undertaken with A549 cells. The values in panels A and D, are the mean ± SEM of triplicate evaluations and the data in panels B, C, E and F are representative of three similar evaluations. In panels C and F blue is DAPI, green is cytoskeleton and red is CHI3L1. (*** p<0.001; ****p<0.0001 by t-test). Scale bar=25μm, it applies to every subpanel of Panels C and F.

### CHI3L1 is produced by cells with mutant *EGFR*

Studies were next undertaken to define the effects of NSCLC-associated *EGFR* mutations on the production of CHI3L1. These experiments demonstrated that HCC827 cells, which have an exon 19 EGFR mutation (29), are potent producers of CHI3L1 mRNA and protein at baseline and that these events are exaggerated by EGF and or TGF-α (Figure 2, A-C). This stimulation was dose-dependent and most prominent after 48-72 hours of incubation with these physiologic ligands of EGFR (Figure 2 and Supplement Figure S2). Similar results were seen with immunohistochemistry which highlighted the accumulation of nuclear and cytoplasmic CHI3L1 in HCC827 cells at baseline and the stimulatory effects of EGFR ligands on HCC827 cell CHI3L1 expression and production (Figure 2, panel C). H1975 cells, which have both the L858R and T790M mutations, had similar effects. (Figure 2, D-F and Supplemental Figure S3). Importantly, cells with NSCLC-associated *EGFR* mutations produced more CHI3L1 than similarly stimulated NHBE cells or transformed cells with wild type EGFR like A549 cells (Supplemental Figure S4). Overall, these studies highlight significant accumulation and production of CHI3L1 in epithelial cells with mutant EGFR.

**Figure 2.**
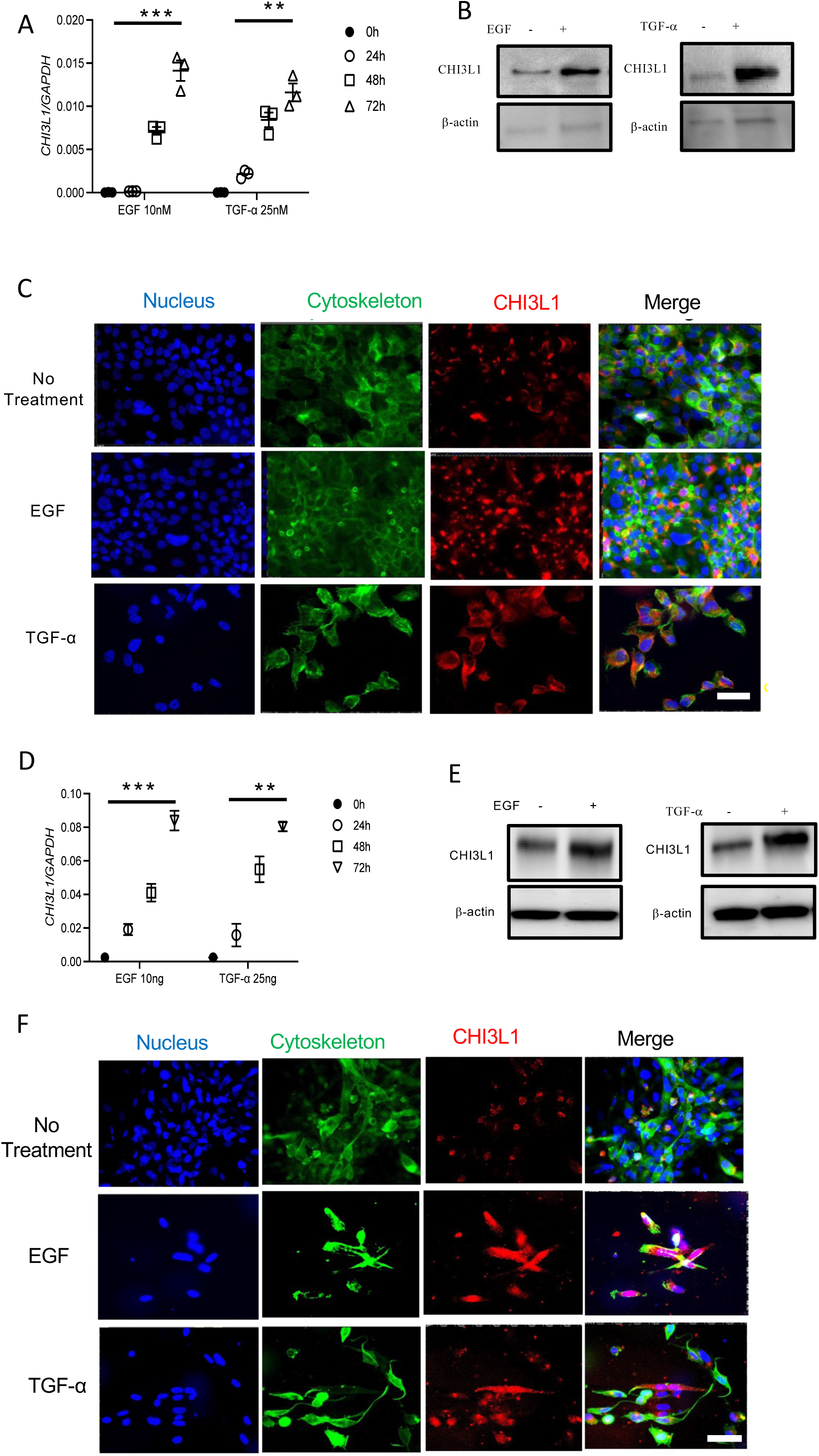
Induction of the expression and production of CHI3L1 in cells with EGFR mutations. HCC827 cells were incubated in the presence or absence of EGF or TGF-α. Triplicate determinations of the levels of mRNA encoding CHI3L1 at the noted time points can be seen in panel A. The accumulation of CHI3L1 protein at 72 hours can be seen with the Western blotting in panel B and double label IF is seen in panel C. In panels D-F, similar evaluations were undertaken with H1975 cells. The values in panels A and D, are the mean ± SEM of triplicate evaluations and the data in panels B, C, E and F are representative of three similar evaluations. In panels C and F blue is DAPI, green is cytoskeleton and red is CHI3L1. (**P<0.1, *** p<0.001 by t-test). Scale bar=25μm, it applies to every subpanel of Panels C and F.

### CHI3L1 forms a positive feedback loop in the EGFR Axis

To further understand the relationships between CHI3L1 and EGFR, we next defined the effects of recombinant (r) CHI3L1 on components of the EGFR axis. These *in vitro* studies demonstrate that rCHI3L1 is a potent, dose-dependent, stimulator of EGFR and its physiologic ligands EGF and TGF-α in cells with mutant EGFR (Figure 3, A-C). These stimulatory effects are CHI3L1-dependent because they were ameliorated by treatment with anti-CHI3L1 (FRG) antibodies (Figure 3, D-I). To determine if similar effects were seen *in vivo*, we compared the levels of mRNA encoding EGFR and its ligands in lungs from transgene negative and lung targeted CHI3L1 transgene positive mice. These studies demonstrate that CHI3L1 is a potent stimulator of EGFR and its ligands EGF and TGF-α in lungs from CHI3L1 transgenic mice (Figure 3, panels J-L). Thus, rCHI3L1 and transgenic CHI3L1 are potent stimulators of EGFR and its ligands in cells with WT and mutant EGFR and lungs from transgenic mice, respectively. When viewed in combination, one can see how the ability of EGFR activation to stimulate the production of CHI3L1 and the ability of CHI3L1 to stimulate EGFR and its ligands sets up a CHI3L1-dependent, positive feedback loop in the EGFR axis.

**Figure 3.**
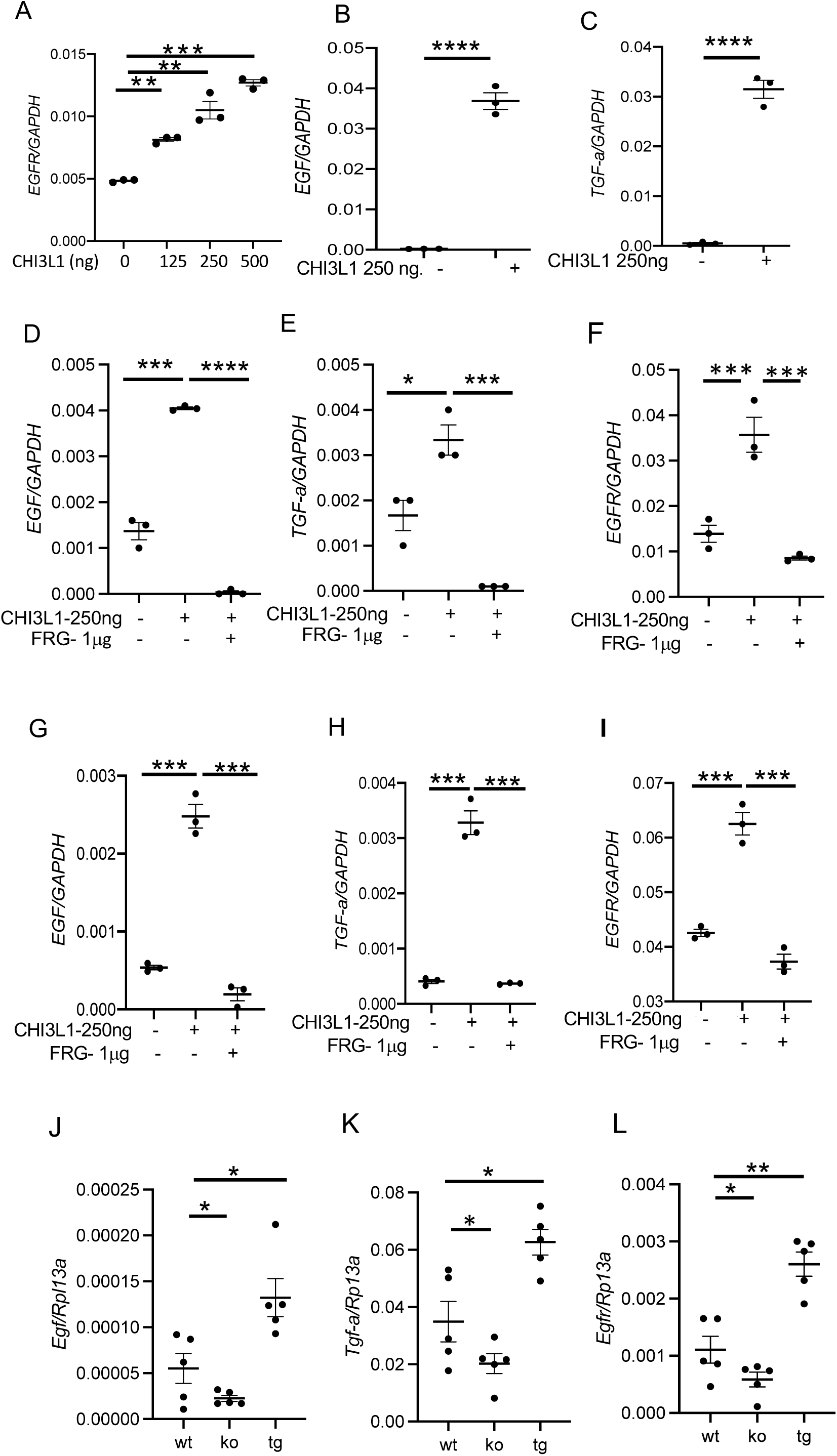
CHI3L1 elicits a positive feedback loop in the EGFR Axis. H1975 cells were incubated with the noted concentration of rCHI3L1 and the levels of mRNA encoding EGFR, EGF and TGF-α were evaluated in panels A-C. The importance of CHI3L1 in these inductive events was assessed in panels D-F by comparing the expression of EGF, TGF-α and EGFR in cells incubated in the presence and absence of anti-CHI3L1 (FRG) antibody. HCC827 cells were similarly incubated with the noted concentrations of rCHI3L1 in the presence and absence of FRG in panels G-I. In panels J-L the levels of mRNA encoding EGFR, EGF and TGF-a were assessed in lungs from wild type (wt) mice, CHI3L1 null mutant mice (ko) and CHI3L1 overexpression transgenic mice (tg). The values in panels A-I represent the mean ± SEM of a minimum of 3 evaluations under each condition. The values in panel J-L represent the mean ± SEM of evaluations in lungs from a minimum of 5 (or more) animals. (*p<0.05; ** p<0.01; *** p<0.001 by One way ANOVA multiple comparisons).

### CHI3L1 is auto-induced in cells with WT and mutant *EGFR*

Additional experiments were undertaken to compare the expression and accumulation of CHI3L1 in cells with WT or mutant *EGFR* that were stimulated with rCHI3L1. These studies demonstrated that CHI3L1 is auto-induced by rCHI3L1 in NHBE cells, cells with WT *EGFR* (A549 cells) and cells with mutant *EGFR*. Similar results were seen in experiments with H1975 cells and HCC827 cells which have differing NSCLC-associated *EGFR* mutations (Figure 4, A-C and Supplemental Figure S4). The studies noted above highlight a CHI3L1-driven positive feedback loop in the EGFR axis. These studies add to our understanding of this feedback loop by highlighting a CHI3L1 auto-induction mechanism that reinforces and feeds this feedback mechanism forward.

**Figure 4.**
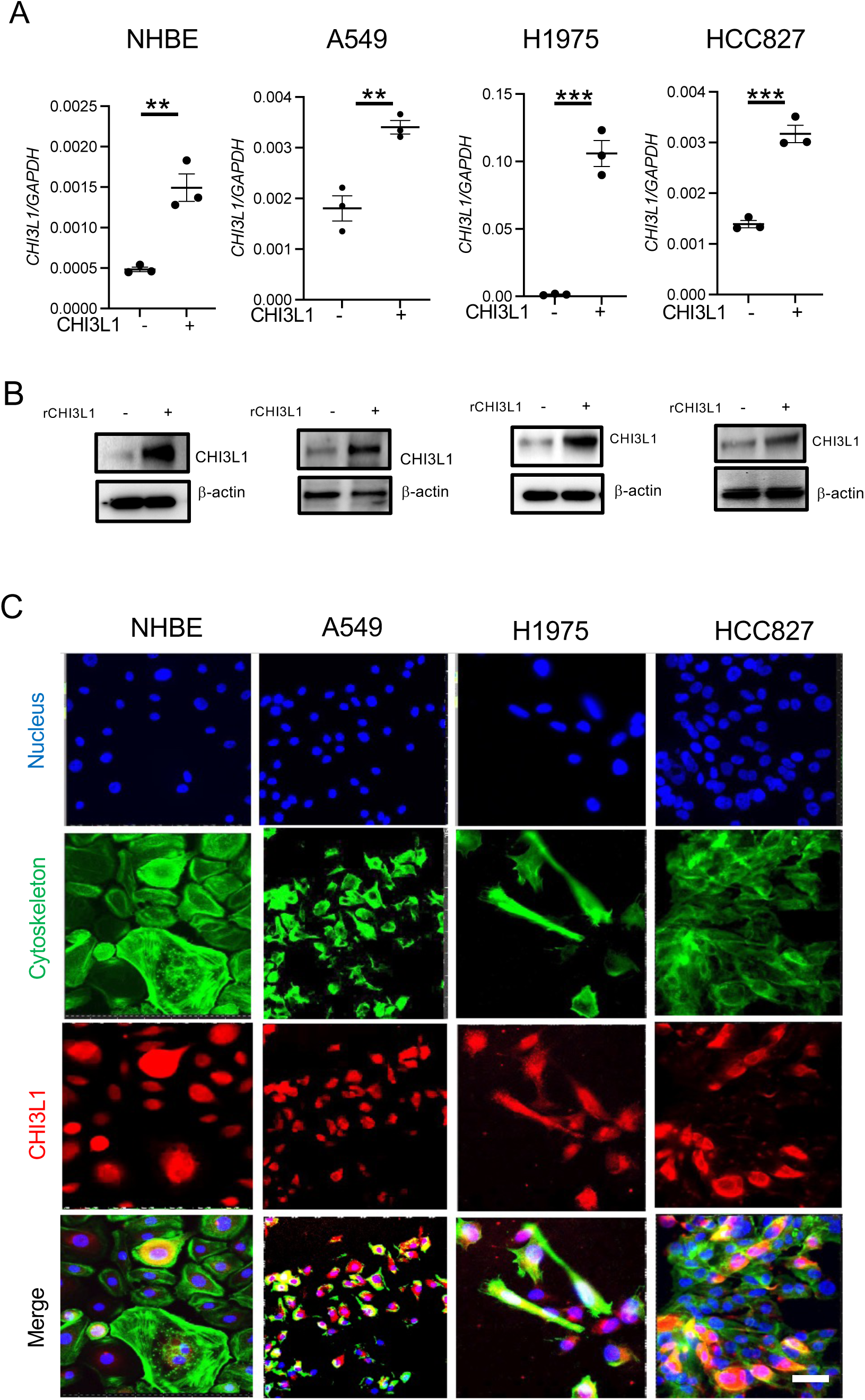
CHI3L1 is auto-induced in cells with WT and mutant EGFR. NHBE, A549, H1975 and HCC827 cells were incubated in the presence and absence of rCHI3L1 and the levels of mRNA encoding CHI3L1 and CHI3L1 protein accumulation were evaluated. The levels of mRNA encoding CHI3L1 were evaluated using qRT-PCR as illustrated in panel A. The levels of CHI3L1 protein accumulation can be seen in the Western blot evaluations and immunocytochemistry in panel B and C. The values in panel A represent the mean ± SEM of at least triplicate evaluations. The values in panels B and C are representative of at least 3 similar evaluations. (*p< 0.05 by t-test). Scale bar=25μm, it applies to every subpanel of Panel C.

### CHI3L1 inhibitors and TKIs, interact in the regulation of the EGFR Axis *In vitro*

TKI have become the first line of treatment for patients with *EGFR* mutant NSCLC. This appreciation started with first generation TKI such as gefitinib or erlotinib (30). Most recently the FLAURA study demonstrated that osimertinib, a 3^rd^ generation TKI, leads to better outcomes than earlier generation TKIs in the first-line therapy for advanced *EGFR*-mutant NSCLC with classical *EGFR* mutations (31). To gain insight into these therapies, we characterized the effects of osimertinib on EGFR activation, CHI3L1 auto-induction and the survival of H1975 and HCC827 cells which have NSCLC-associated *EGFR* mutations. These studies demonstrated that, individually, osimertinib is a modest inhibitor and that anti-CHI3L1 (FRG) is a more potent inhibitor of CHI3L1 auto-induction and CHI3L1 stimulation of EGFR and EGF in these HC8227 cells (Figure 5A-C). Interestingly, when osimertinib and FRG were used in combination impressive interactions were appreciated which were at least additive in nature (Figure 5, A-C). Impressive synergistic inhibition of CHI3L1 autoinduction and EGF and EGFR expression were also seen when lower doses of osimertinib and FRG were employed (Figure 5, D-E). In these experiments, lower doses of osimertinib (1nM) and FRG (5ng) that individually did not cause significant alterations in CHI3L1 autoinduction or EGF induction were impressive inhibitors of CHI3L1 auto-induction and EGF when used in combination (Figure 5D-E).

**Figure 5.**
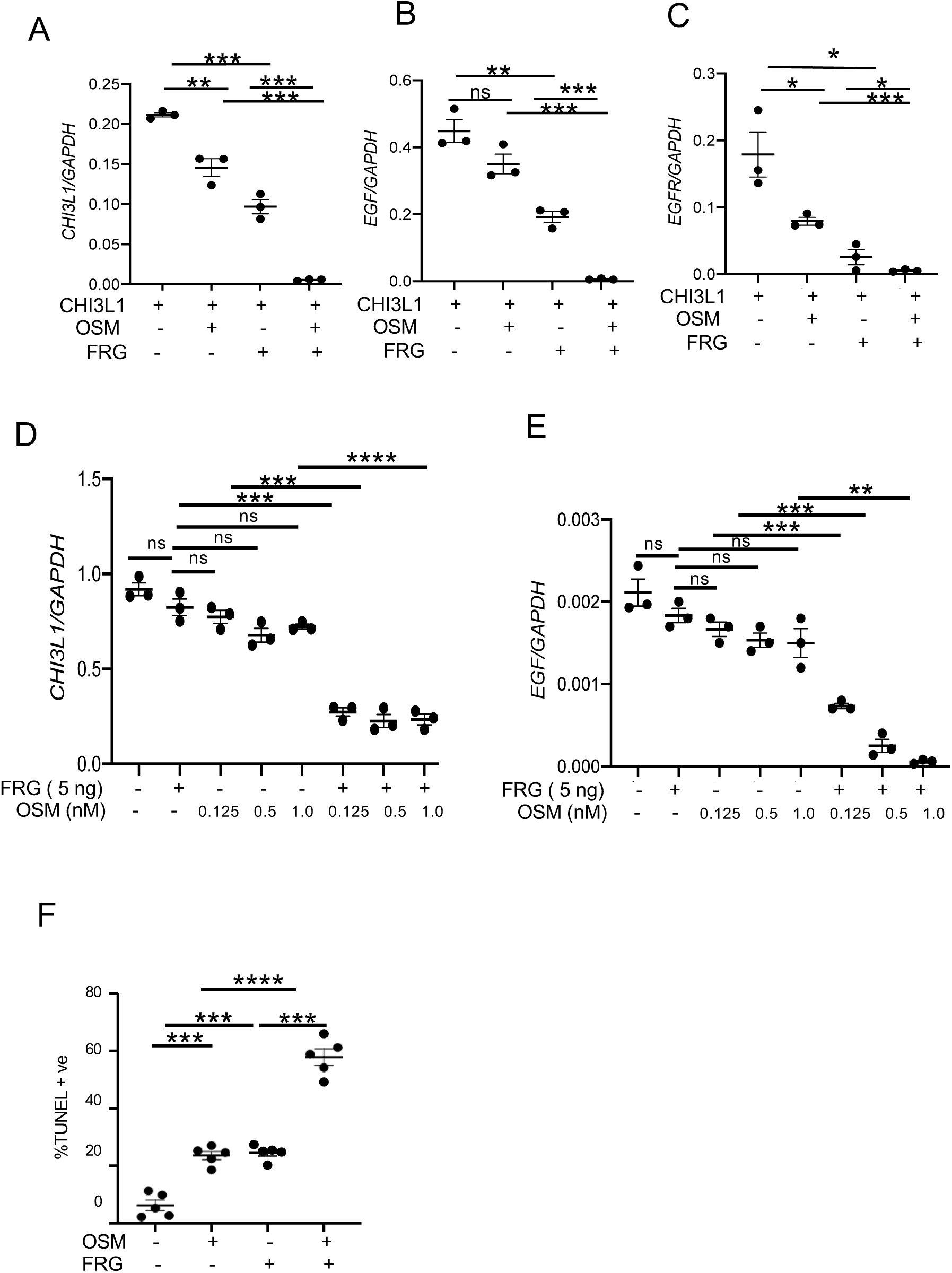
Effects of CHI3L1 inhibitors and TKI, alone and in combination, on the EGFR axis *in vitro*. HCC827 cells were incubated with rCHI3L1 and the effects of osimertinib (OSM) and FRG on the expression of CHI3L1, EGF and EGFR were evaluated by qRT-PCR (panels A-C). In panels D and E, H1975 cells were incubated with low doses of osimertinib (OSM) and FRG, alone or in combination, and the levels of mRNA encoding CHI3L1 and EGF were evaluated. In panel F, H1975 cells were incubated with osimertinib (OSM) and FRG antibody, alone or in combination, and the levels of apoptosis were evaluated by TUNEL staining. (*p<0.05; **p<0.01; *** p<0.001, ****p<0.0001 by ANOVA with multiple comparisons). ns, not significant.

To see if apoptotic cell death was involved in the combinatorial effects of gefitinib and FRG, alone or in combination, TUNEL evaluations of DNA cell injury were employed. In these experiments gefitinib and FRG individually caused cell apoptosis. Interestingly, when gefitinib and FRG were used in combination, impressive increases in cellular apoptosis were seen. (Supplemental Figure S5). Similar effects on apoptosis were seen with osimertinib and FRG, alone or in combination (Figure 5F). These studies demonstrate that TKI and anti-CHI3L1 (FRG) individually, and more potently in combination, inhibit CHI3L1 auto-induction and EGFR activation while inducing cellular apoptosis.

### Anti-CHI3L1 (FRG) reverses therapeutic resistance to osimertinib

A major rate limiting step in the treatment of *EGFR* mutant NSCLC is the development of resistance to the therapeutic agents. To gain insight, into the relationship between this resistance and CHI3L1, we grew HCC827 cells in osimertinib until therapeutic resistance was acquired (32). We then compared the gene expression and survival of these HCC8270R2 cells when incubated in the presence and absence of anti-CHI3L1 (FRG). As can be seen in Figure 6, osimertinib and FRG individually caused modest decreases in CHI3L1 auto-induction and the expression of CHI3L1, EGF and EGFR (Figure 6A). Importantly, the addition of FRG to cells in osimertinib elicited an impressive synergistic inhibition of the expression of CHI3L1, EGF, and EGFR (Figure 6A). These effects were associated with similar changes in cell survival with osimertinib and FRG individually causing prudent increases in cellular apoptosis and the two therapeutic agents in combination causing an impressive synergistic increase in cell apoptosis (Figure 6B). Kelch like ECH associated protein 1 (*KEAP1*) transcriptional activity has been associated with EGFR activation (33) . In accord with the ability of the tumor suppressor gene *KEAP1* to modulate TKI sensitivity (34), these experiments were associated with the opposite alterations in *KEAP1* with osimertinib and FRG individually causing modest and osimertinib and FRG in combination causing increases in *KEAP1* expression and protein accumulation (Figure 6, C and D). Similar results were seen with other tumor suppressor genes including retinoblastoma (*RB*1), *P53* and *PTEN* (Figure 6E). These studies demonstrate that CHI3L1 inhibition reverses osimertinib resistance by inhibiting CHI3L1 auto-induction and inhibiting the expression of EGFR and its ligands while inducing apoptosis and stimulating production of KEAP1 and other tumor suppressors.

**Figure 6.**
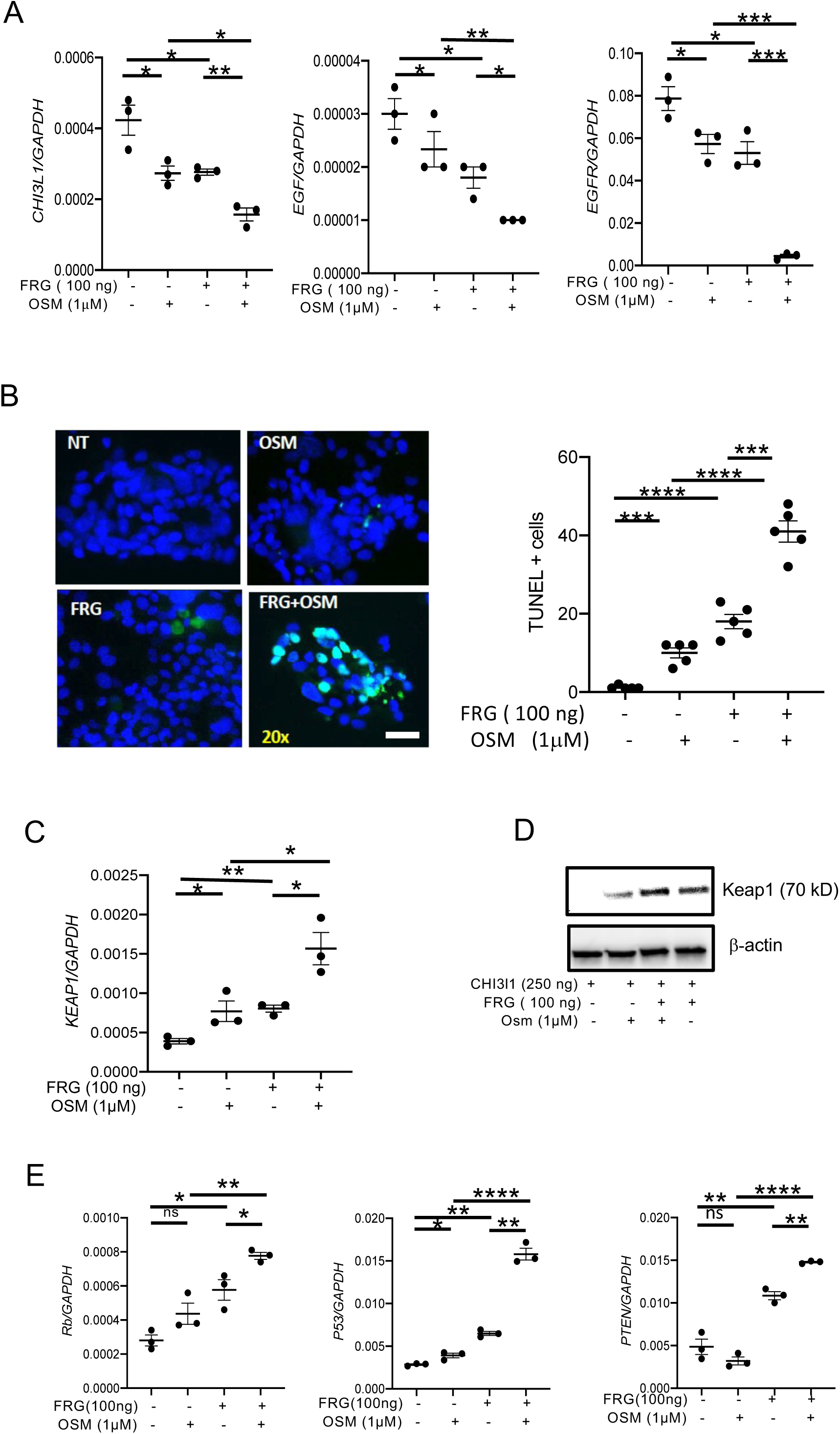
Anti-CHI3l1 (FRG) reverses therapeutic resistance to osimertinib and regulates tumor suppressor genes. HCC827 cells were chronically incubated with osimertinib to develop osimertinib resistant HCC8270R2 cells. The effects of CHI3L1 inhibitors and TKI, alone or in combination, were then evaluated by incubating the HCC8270R2 cells with osimertinib, alone or in combination with FRG and evaluating the levels of mRNA encoding CHI3L1, EGF and EGFR (panel A). The ability of osimertinib to induce HCC8270R2 cell death was also evaluated. In this experiment HCC8270R2 cells were incubated with osimertinib and FRG, alone or in combination, and TUNEL evaluations were undertaken (Panel B). In these experiments, studies were also undertaken to determine if osimertinib and FRG, alone or in combination altered the expression and or accumulation of tumor suppressor genes. The effects on *KEAP1* mRNA and protein can be seen in panels C and D respectively. The effects on *RB1*, *P53* and *PTEN* are illustrated in panel E. (* p<0.05, **p<0.01; *** p<0.001, ****p<0.0001 by ANOVA with multiple comparisons). ns, not significant. Scale bar=25μm, it applies to every subpanel of Panel B.

### CHI3L1 inhibitors and TKI, alone and in combination, inhibit pulmonary metastasis *in vivo*

As noted above, malignancies with *EGFR* driver mutations frequently initially respond to and then fail to respond and develop metastasis as they develop TKI resistance. Thus, to further our understanding of TKI resistance and the relationship between FRG and TKI, we characterized the effects on pulmonary metastasis of TKI and FRG, alone and in combination in B16 cell challenged mice. When used individually, FRG (intraperitoneally), and gefitinib or osimertinib (orally) inhibited B16-F10 pulmonary metastasis (Figure 7, A and B). When used in combination, at selected doses, impressive synergistic inhibition of tumor metastasis was seen (Figure 7, A and B). These studies demonstrate that anti-CHI3L1 and TKI, individually and more potently in combination, inhibit pulmonary metastasis.

**Figure 7.**
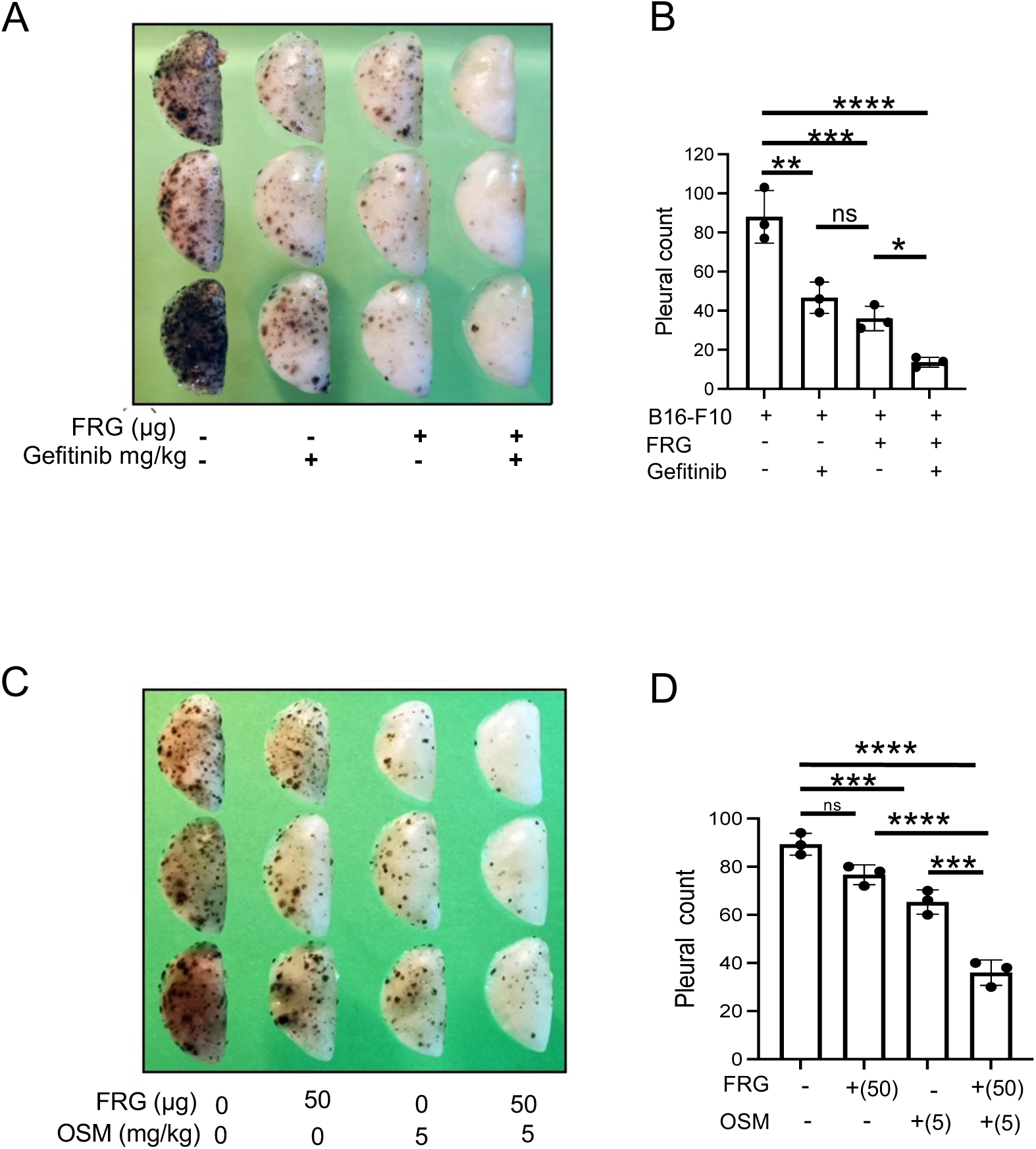
Anti-CHI3L1 and TKI interact to inhibit tumor metastasis. WT mice B16-F10 were challenged with (B16) melanoma cells or control vehicle and treated with control IgG, FRG and or TKIs, alone or in combination. Melanoma lung metastasis was evaluated 2 weeks later. (A) Representative lungs from mice treated with control IgG, FRG and or gefitinib, alone or in combination. (B) The number of pleural melanoma colonies was quantitated in the lungs from the mice in panel A. (C) Representative lungs from mice treated with IgG, FRG and osimertinib, alone and in combination. (D) The number of pleural melanoma colonies was quantitated in the lungs from the mice in panel C. In panels A and C, each dot is representative of an individual animal. Panels A and C are representative of at least 3 evaluations. The values in panels B and D represent the mean ± SEM of the evaluations represented by the individual dots in the lungs from the experiments illustrated in panel A and C respectively. \**P*<0.05. \*\*\**P*<0.001; \*\*\*\**P*<0.0001. by ANOVA with multiple comparisons). ns, not significant.

## Discussion

Aberrant activation of EGFR by erythroblastic leukemia viral homologue (*ERBB*) ligands contributes to the pathogenesis of a variety of cancers including NSCLC (5). In these tumors, EGFR and its ligands are stimulated and activated resulting in cancer development and progression(5). To determine if CHI3L1 contributes to the stimulation of and activation of EGFR and its ligands, studies were undertaken to define the relationships between CHI3L1 and the EGFR axis. These studies demonstrate, for the first time, that EGFR activation by its physiologic ligands or as a result of activating mutations stimulates the production and activation of CHI3L1. They also demonstrate, for the first time, that CHI3L1 feeds back to form an amplifying feedback loop that stimulates the production and expression of EGFR and its ligands in cells with WT and mutant *EGFR*. This is in accord with the known importance of autocrine, paracrine and juxtacrine EGFR-based feedback loops that activate intracellular signaling in the original cell and in neighboring and distant cells (5, 35). Lastly, they also highlight the autoinduction of CHI3L1 with CHI3L1 stimulating EGFR and its ligands which, in turn, further stimulate the production of CHI3L1. This stimulation of CHI3L1 was mediated by EGFR because it was abrogated by treatment with TKI. It was also seen in cells with WT *EGFR* and common *EGFR* mutations (L858R, exon 19 deletion) and less common EGFR mutations (G719X, L861X, Exon19 Insertion). When viewed in combination, these studies highlight a newly appreciated intimate relationship between CHI3L1, EGFR and its ligands with each stimulating the production of the other to augment tumor growth and progression.

The discovery that the EGFR axis controls tyrosine kinase activation led to concerted efforts to develop tyrosine kinase inhibitors (TKI) as therapeutics for patients with *EGFR* mutant NSCLC. As a consequence, there are three generations of approved TKI therapeutics which have changed our therapeutic approach to NSCLC (6). All three generations of TKIs have demonstrated higher objective response rates (ORR) and prolonged progression free survivals (PFS) compared to standard chemotherapy (6, 30, 36–38). As a result, TKIs have become the first-choice therapeutics for patients with advanced *EGFR* mutant NSCLC. Approximately, 20-30% of patients with *EGFR* mutant NSCLC exhibit primary resistance to EGFR TKIs. In contrast, the majority of the remaining patients exhibit responses to these interventions (6, 30). These responses, however, can be heterogeneous in terms of depth and duration. Importantly, despite the remarkably high objective response rates to TKIs, the majority of these patients develop therapeutic resistance with progression free survival (PFS) ranging from 9-18 months (8, 39). Our studies provide new insights into the mechanisms that contribute to these effects of the TKIs. Specifically, they demonstrate, for the first time, that treatment with TKIs inhibit the induction of CHI3L1 that is caused by EGFR activation by its physiologic ligands or oncogenic *EGFR* mutations. They also demonstrate that TKIs abrogate the ability of CHI3L1 to feedback to stimulate EGFR and its ligands and contribute to CHI3L1 autoinduction.

In keeping with the high degree of tumor heterogeneity and adaptive cellular signaling, a variety of mechanisms have been shown to contribute to TKI therapeutic resistance in the treatment of NSCLC (8). These resistance mechanisms can exist at the time of diagnosis when initial treatment is being initiated. They also occur during TKI treatment or after the failure of treatment with other EGFR inhibitors (8). In the latter, EGFR-TKI resistance mechanisms can be broadly grouped into EGFR-dependent and EGFR-independent pathways (8). The most common resistance mechanism results from the development of the so called “gatekeeper” T790M mutation which changes the affinity of first and second generation TKI to the ATP binding site of EGFR (8, 40). This mutation, however, has been overcome by the 3^rd^ generation TKI osimertinib which has been approved by the FDA for the treatment of patients with EGFR mutant NSCLC who have acquired this mutation after progression on prior TKI therapy (8). Our studies demonstrate that CHI3L1 contributes to TKI resistance in cells treated with first generation drug gefitinib and third generation drug osimertinib. They also demonstrate that CHI3L1 contributes to TKI resistance in cells with WT EGFR, common and rare EGFR mutations and cells with T790M mutations. Importantly, they also demonstrate that interventions that inhibit CHI3L1 abrogate the induction of EGFR and its ligands and the autoinduction of CHI3L1 thereby inhibiting the activation of the EGFR axis via EGFR-dependent and -independent mechanisms.

To further understand the mechanisms that contribute to TKI resistance in *EGFR* mutant NSCLC, we chronically exposed HCC827 cells (exon 19 deletion) to osimertinib. When osimertinib treatment was first initiated it induced tumor cell death and decreased tumor cell proliferation. However, osimertinib resistance eventually developed with osimertinib-induced cell death and osimertinib inhibition of cell proliferation being abrogated (32). The role of CHI3L1 in this TKI resistance was then evaluated by determining if the anti-CHI3L1 (FRG) was effective in reversing the osimertinib resistance in these cells. The osimertinib-resistant cells did not manifest significant TUNEL staining when incubated with osimertinib at baseline. They did, however, contain exaggerated levels of mRNA that encoded EGFR and its ligands. In contrast, the addition of FRG caused an impressive increase in TUNEL staining and cell apoptosis. FRG also inhibited the ability of CHI3L1 to stimulate EGFR and its ligands and inhibited CHI3L1 autoinduction. These observations are in accord with prior observations from our laboratory and others which demonstrated that CHI3L1 is an impressive inhibitor of apoptosis which is mediated, at least in part, by its ability to activate MAPK and AKT anti-apoptotic signaling pathways (41–44).

Tumors are frequently classified based on their driver mutations while the function of coincident alterations in tumor suppressor genes are rarely considered (34). However, there is emerging evidence that the interplay between oncogenic drivers and tumor suppressors can influence tumor fitness and alter therapeutic outcomes (34). In lung adenocarcinomas, *EGFR* is one of the most frequently mutated driver genes where it occurs on a background of diverse tumor suppressor gene alterations (34, 45–49). Genomic alterations in tumor suppressor genes such as *RB1* and *P53* correlate with outcomes in TKI therapy (34, 48, 50, 51). Inhibition of several tumor suppressor genes has been demonstrated to result in larger tumors in murine models of *EGFR* mutant NSCLC (34). Studies have also defined a link between loss of specific tumor suppressor genes and responses to osimertinib in *EGFR* mutant lung cancer (34). However, the contribution of tumor suppressor genes to TKI sensitivity and drug resistance is only now beginning to be understood (34). To address this issue, we evaluated the expression of *P53*, *KEAP1*, *PTEN* and *RB1* in osimertinib resistant HCC827OR2 cells treated with FRG or its isotype control. These studies demonstrate that FRG stimulates the expression of these tumor suppressors while restoring the osimertinib sensitivity of these cells. These findings are in accord with prior studies that demonstrate that CHI3L1 ubiquitinates p53 to target it for degradation (52), studies from our laboratory that demonstrate that CHI3L1 inhibits *PTEN* (26, 28) and studies that demonstrate that RB1 inactivation is a major driver of tumor growth in *EGFR* mutant cancers (34). They are also in accord with studies that demonstrate that *KEAP1* is a negative regulator of the transcription factor NRF2 which alters cell metabolism and cellular antioxidant responses (34). *KEAP1* inactivation has also been shown to decrease tumor cell sensitivity to osimertinib (34). Patients with tumors harboring mutations in *KEAP1* axis genes have a significantly shorter time to treatment discontinuation compared to those with *KEAP1* wild-type tumors (34, 50, 53).

Resistance to TKI therapeutics can manifest with tumor progression and metastasis. Because the inhibition of CHI3L1 can restore the sensitivity of *EGFR* mutant NSCLC cells to osimertinib, studies were also undertaken to determine if anti-CHI3L1 (FRG) could interact with TKI (gefitinib or osimertinib) in the regulation of cancer metastasis. These studies highlight impressive interactions between these moieties with FRG interacting in a synergistic manner with gefitinib or appropriate doses of osimertinib to inhibit lung metastasis. Although it is well known that targeted therapeutics like TKI can be quite effective when initially administered, it is also known that these responses are often not durable (54). It is also appreciated that the blockade of one tumorigenic pathway can lead to compensatory activation of several others (55, 56). As a result, many have turned to combination therapeutics to achieve enhanced anti-tumor responses (54). Our studies highlight the enhanced antitumor responses that are seen with the combination of FRG and gefitinib and the combination of FRG and osimertinib. These observations suggest that these combinations of agents may be particularly useful in the treatment of NSCLC and other metastatic malignancies. Additional investigation will be required to further define the optimal parameters of the use of this combination therapeutic approach.

The studies in this manuscript focus predominantly on the role(s) of and interactions between the CHI3L1 and the EGFR axis in NSCLC. They demonstrate that these regulator pathways induce and activate each other and form a positive (+ve) feedback loop between EGFR, its physiologic ligands and CHI3L1. They also demonstrate, for the first time, that CHI3L1 plays an important role in the development of resistance to therapeutic TKI, that anti-CHI3L1 interacts in a synergistic manner with osimertinib to suppress tumor metastasis and that anti-CHI3L1 reverses osimertinib therapeutic resistance. It is important to point out that EGFR mutations are also seen in and play important role(s) in the pathogenesis of other tumors including glioblastoma and colorectal malignancies (5). A particularly interesting association can be seen in glioblastoma multiforme (GBM) (57). CHI3L1 is prominently expressed in these tumors where it predicts radiation resistance and a poor prognosis (57). More recent investigations from our laboratory and others, also demonstrated that CHI3L1 drives mesenchymal differentiation of GBM stem cells (58) and that EGFR amplification is also a prominent feature of GBM. Based on these findings it is tempting to speculate that interactions such as the ones described above in NSCLC are operative in these other disorders. It is also tempting to speculate that appropriately humanized antiCHI3L1 antibodies can be useful, alone or in combination with TKI, in the treatment of these other disorders. Additional, investigation will be required to test the veracity and utility of these hypotheses.

In conclusion, our studies demonstrate that CHI3L1 plays a number of important roles in the pathogenesis of *EGFR* mutant NSCLC and malignant metastasis. These studies highlight the ability of EGFR activation by its physiologic ligands or activating *EGFR* mutations to stimulate the production of CHI3L1 and the ability of CHI3L1 to feedback to form an amplifying loop that stimulates EGFR and its ligands and further stimulates CHI3L1 via EGFR-dependent mechanisms. They also demonstrate that the resistance to TKI that limits the efficacy of therapeutics like osimertinib can be reversed with anti-CHI3L1 and highlight the importance of EGFR-dependent and EGFR-independent pathways and tumor suppressors including *P53*, *KEAP1*, *PTEN*, and *RB1* in this response. Lastly, they highlight the ability of TKIs and FRG to interact in a synergistic manner to inhibit tumor metastasis. When viewed in combination, these studies strongly support the contention that CHI3L1 plays a critical role in the pathogenesis of and is a critical therapeutic target in *EGFR* mutant NSCLC. They also demonstrate that interventions that inhibit CHI3L1 can augment the therapeutic efficacy of targeted EGFR TKI therapeutics in NSCLC and malignant metastasis. Additional investigations of the biology of CHI3L1 in the EGFR axis and the utility of interventions that target CHI3L1 in patients with *EGFR* mutant and other cancers are warranted.

## Materials and Methods

### Cell line

Normal lung epithelial cells and lung cancer cell lines were purchased from the ATCC: NHBE (PCS-300-010^™^), A549 (CRM-CCL-185), H1975 (CRL-5908) and HCC827 (CRL-2868). Osimertinib-resistant HCC827OR2 cells were established according to the published protocol (59). These osimertinib cells were maintained in a growth medium containing osimertinib (1µM).

### Genetically modified mice

Lung-specific CHI3L1 overexpressing transgenic mice in which CHI3L1 was targeted to the lung with the CC10 promoter (*CHI3L1* Tg) and CHI3L1 null mutant mice (*Chil1*^-/-^) have been generated and characterized by our laboratory as previously described (11, 60). These mice were between 6-12 weeks old when used in these studies. All animals were humanely anesthetized with Ketamine/Xylazine (100mg/10mg/kg/mouse) before any intervention. The protocols that were used in these studies were evaluated and approved by the Institutional Animal Care and Use Committee (IACUC) at Brown University.

### Western blot analysis

Protein lysates from cells were prepared with RIPA lysis buffer (ThermoFisher Scientific, Waltham, MA, USA) containing protease inhibitor cocktail (ThermoFisher Scientific) as per the manufacturer’s instructions. 20 to 30 µg of lysate protein was subjected to SDS-PAGE electrophoresis on a 4–15% gradient mini-Protean TGX gel (Bio-rad, Hercules, CA, USA). It was then transferred to a membrane using a semidry method with a Trans-Blot Turbo Transfer System (Bio-Rad). Membranes were blocked with Tris-buffered saline with Tween20 (TBST) with 5% skimmed milk for 1 hour at room temperature. After blocking, the membranes were incubated with the primary antibodies overnight at 4°C in TBST and 3% BSA. The primary antibodies used in this study are anti-(α) human CHI3L1 (R&D. AF2599-SP) and α-KEAP-1(Santa Cruz Biotechnology. 8047), β-actin (Santa Cruz Biotechnology. Sc47778) antibodies. The membranes were then washed 3 times with TBST and incubated with secondary antibodies in TBST, 5% skimmed milk for 1 hour at room temperature. After 3 additional TBST washes, Supersignal West Femto Maximum Sensitivity Substrate Solution (Thermofisher Scientific) was added on to the membrane and immunoreactive bands were detected by using a ChemiDoc (Bio-Rad) imaging system.

### RNA extraction and Real-time qPCR

Total cellular RNA was obtained using TRIzol reagent (ThermoFisher Scientific) followed by RNA extraction using RNeasy Mini Kit (Qiagen, Germantown, MD) according to the manufacturer’s instructions. mRNA was measured and used for real time (RT)-PCR as described previously (11, 19). The primer sequences used in these studies are summarized in supplemental Table S1. Ct values of the test genes were normalized to the internal housekeeping gene *GAPDH* or *Rpl13a*.

### Measurement of cellular apoptosis and cytotoxic cell death responses

TUNEL staining using fluorescein-labeled dUTP was employed to assess DNA injury, apoptosis and cytotoxic cell death responses. After 72 hours incubation with test and control antibodies, cells were fixed in the 4% paraformaldehyde in PBS, permeabilized and blocked. In between each step, the cells were washed twice with PBS (1X). The cells were then stained with the *in-situ* cell death detection kit, fluorescein (Roche, Mannheim, Germany) as per the manufacturer’s instructions.

### Immunofluorescence staining

To evaluate expression of CHI3L1, immunofluorescence (IF) staining was carried out using antibodies against CHI3L1 (FRG antibody). After 72 hours incubation period with test and control antibodies, cells were fixed in the 4% paraformaldehyde in PBS, permeabilized and blocked. Then the cells were incubated overnight at 4°C with primary antibodies noted above, washed twice with PBS (1X) and incubated for 2 hours at 37°C with secondary detection antibodies (1:500) and phalloidin (1: 3000) (Invitrogen, Waltham, MA) for cytoskeleton staining. The cells were then washed, mounted with VECTASHIELD® antifade mounting medium (Vector Laboratories Inc.) and evaluated at 20X via fluorescence microscopy.

### Melanoma lung metastasis and antibody treatment

B16-F10, a mouse melanoma cell line, was purchased from ATCC (Cat#: CRL-6475, Manassas, VA) and maintained in Dulbecco’s Modified Eagles Medium (DMEM) supplemented with 10% Fetal bovine Serum (FBS) and 1% penicillin & streptomycin. When the cells formed an 80% confluent monolayer, they were collected, adjusted to the concentration of 10^6^ cells/ml and injected into the mice via their lateral tail veins (2×10^5^ cells/mouse in 200μl of DMEM). As previously described (48), intraperitoneal injection of the noted doses of anti-Chi3l1 (FRG) antibody (50 and 200 µg/ mouse) and different doses of gefitinib (1mg/Kg) and Osimertinib (5mg/Kg), alone and in combination and isotype control IgG (IgG2b) were started on the day of the B16 tumor cell challenge and continued every other day for 2 weeks. Metastasis was assessed and quantified by counting the melanoma colonies (black dots) on the pleural surface as previously described (26, 61).

### Quantification and Statistical analysis

Statistical evaluations were undertaken with GraphPad Prism software. As appropriate, groups were compared with 2-tailed Student’s *t* test. Values are expressed as mean ± SEM. One way ANOVA test were used for multiple group comparisons. Statistical significance was defined as a level of *P* < 0.05.

## Competing Interests

JAE is a cofounder of Elkurt Therapeutics and is a founder of and stockholder of, and serves on the Scientific Advisory Board for Ocean Biomedical, Inc., which develops inhibitors of 18 glycosyl hydrolases as therapeutics. CGL, CML, BM, and SK serves on consultants for Ocean Biomedical. Inc. JAE, CGL and SK have composition of matter and use patents relating to antibodies against CHI3L1. The other Brown University authors have declared that no conflict of interest exists. K. Politi reports grants and consulting fees from AstraZeneca; grants from Roche/Genentech, D2G Oncology and Boehringer Ingelheim; and consulting fees from Jannssen; and a patent for EGFR^T790M^ mutation testing issued, licensed, and with royalties paid from Molecular Diagnostics/Memorial Sloan Kettering Cancer Center.

