## Supplemental Table and Figures for "Chitinase 3-like-1 (CHI3L1) in the Pathogenesis of Epidermal Growth Factor Receptor Mutant Non-Small Cell Lung Cancer"

**Table.S1**

| Human Gene | Forward primer | Ntd. | Reverse primer | Ntd. |
|---|---|---|---|---|
| GAPDH | GTCTCCTCTGACTTCAACAGCG | 22 | ACCACCCTGTTGCTGTAGCCAA | 22 |
| CHI3L1 | CCACAGTCCATAGAATCCTCGG | 22 | TGCCTGTCCTTCAGGTACTGCA | 22 |
| EGFR | AACACCCTGGTCTGGAAGTACG | 22 | TCGTTGGACAGCCTTCAAGACC | 22 |
| EGF | TTGGTGATGGGAGGATGACT | 20 | GGCCAGTGACTCAGCAGAAA | 20 |
| TGF-a | CTCAGACACATGCCTACACCT | 21 | GAACATCTCTGCTCGTCGCT | 20 |
| KEAP-1 | AGCGACGGTTCTACGTCCA | 19 | GCGACTGTCGGAAGTAGCC | 19 |
| P53 | CCTCAGCATCTTATCCGAGTGG | 22 | TGGATGGTGGTACAGTCAGAGC | 22 |
| Rb1 | AACATGCCCGACCCAATTACA | 21 | GTGTTTCTGCATACCTCATGG | 21 |
| PTEN | TGAGTTCCCTCAGCCGTTACCT | 22 | GAGGTTTCCTCTGGTCCTGGTA | 22 |

| Mouse Gene | Forward primer | Ntd. | Reverse primer | Ntd. |
|---|---|---|---|---|
| *Rpl13a* | AGGGGCAGGTTCTGGTATTG | 20 | TGTTGATGCCTTCACAGCGT | 20 |
| *Chi3l1* | GCTTTGCCAACATCAGCAGCGA | 22 | AGGAGGGTCTTCAGGTTGGTGT | 22 |
| *Egfr* | GGACTGTGTCTCCTGCCAGAAT | 22 | GGCAGACATTCTGGATGGCACT | 22 |
| *Egf* | ACTGGTGTGACACCAAGAGGTC | 22 | CCACAGGTGATCCTCAAACACG | 22 |
| *Tgf-*$\alpha$ | TGATACGCCTGAGTGGCTGTCT | 22 | CACAAGAGCAGTGAGCGCTGAA | 22 |

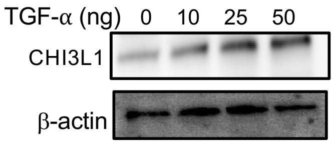


**Figure S1**. Dose response of TGF-α (10, 50 and 100 ng/ml) stimulation of CHI3L1 protein production by NHBE cells evaluated by Western analysis.

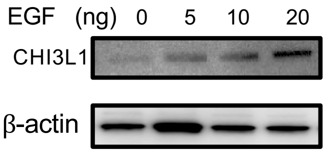


**Figure S2**. Dose response of EGF (5,10 and 20 ng/ml) stimulation of CHI3L1 protein production in HCC827 cells evaluated by Western analysis..

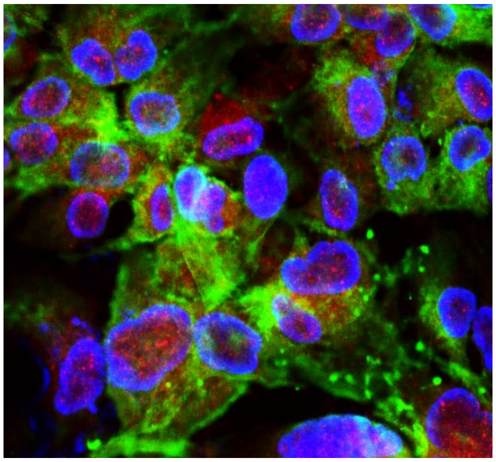

**Figure. S3**. Immunohistochemical evaluation of baseline CHI3L1 (red) in nuclei and cytoplasm in HCC827 cells (DAPI=blue, cytoskeleton: green, CHI3L1:red) 20x.

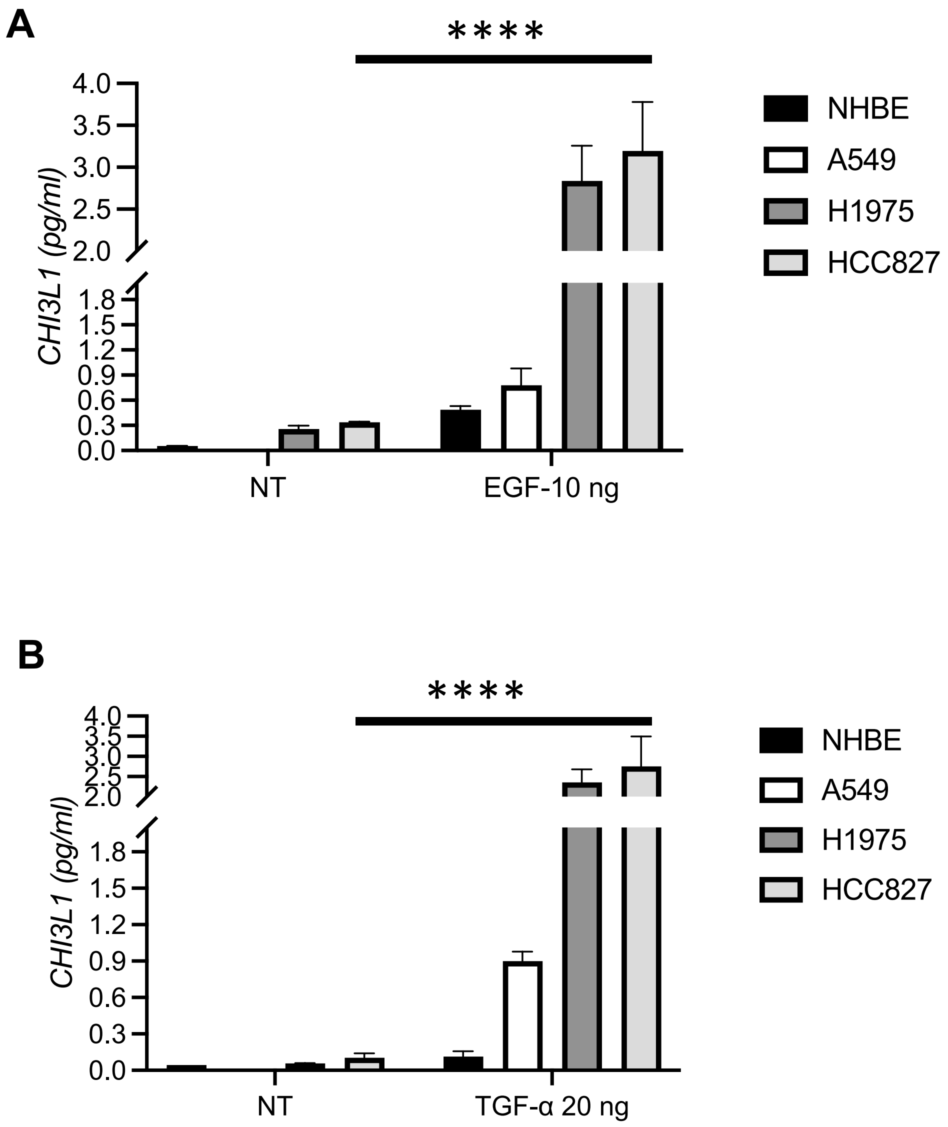


**Figure S4**. ELISA-based comparison of the levels of CHI3L1 secreted by (A) EGF and (B) TGF-$\alpha$ stimulated NHBE, A549, H1975 and HCC827 cells incubated for 72 hours. (*P<0.001) (NT= no treatment)

**A**

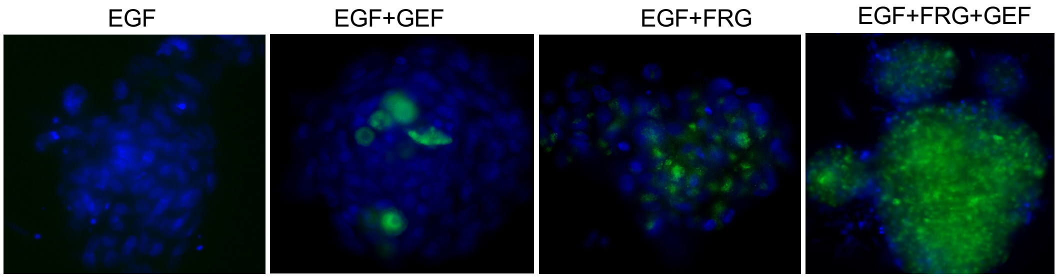


**B**

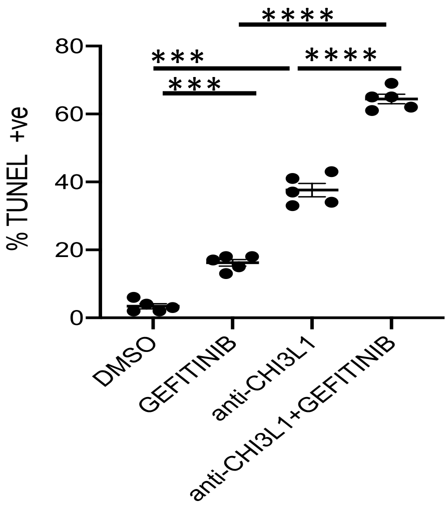


**Figure S5**. TUNEL evaluation of H1975 cells incubated with FRG antibody and Gefitinib, alone and in combination. EGF (20ng/ml) and Gefitinib (100 ng/ml)
